# Diversity of Insecticidal PA1b Homologs among Legume Seeds from Middle Eastern Region

**DOI:** 10.1101/2023.12.04.569987

**Authors:** F. Diya, H. Charles, A. Vallier, L. Karaki, L. Kfoury, P. Da Silva, F. Rizk

## Abstract

Legumes play a central role in various food systems, with significant socio-economic and environmental impacts. Their high protein content, composed mainly of globulins and albumins, makes them valuable for human food and animal feed. Among the albumins, is Pea Albumin 1 b (PA1b), a 37 amino acid peptide, extracted from the seeds of the pea *Pisum sativum*. The protein displays the knottin scaffold and exhibits potent insecticidal activity against certain insects including cereal weevils and mosquitoes. This toxicity is attributed to the coexistence of several isoforms in peas. The natural diversity of PA1b-like molecules within the legume species of the Fabaceae family has been studied using various molecular, biochemical, and bioinformatic tools. Several *A1* genes coding for this peptide have been characterized in soybeans, bean, barrel medick and other legume species. The aim of is study is to precisely characterize partial A1 genes in legumes of the Faboideae subfamily from the Middle East region using PCR homology. Specifically, the research focuses on the sequence structure of Pea Albumin 1 b (PA1b) variants and establishes phylogenetic relationships between these sequences and publicly available A1b homologs. The toxic effects of seed flour containing PA1b-like molecules are assessed, demonstrating that the newly characterized PA1b homologs retain structural conservation. The study observes both conservation and diversification among A1b homologs, consistent with the divergence of lineages within the Fabaceae family. The toxic effects associated with putative A1b molecules are found across different species and within the same species from different geographical origins. In particular, novel candidates such as *Vicia sativa* and *Medicago minima* show promising insecticidal A1b activity. Further analysis of isoforms from these species, including an examination of their expression in different tissues and organs should be undertaken to facilitate the potential use of A1b molecules in agricultural practice.

## Introduction

The Fabaceae family is the third-largest family of angiosperms, with its 800 genera and about 20,000 species^1^. Species in this family are also known as grain legumes, and some are weeds of cereal crops^2^. Legumes play an important role in various food systems, socio-economic and environmental aspects^3^. They are characterized by their high protein content, that complements that of cereals, making them suitable for both human and animal consumption^3,4^. One of their notable features is their ability to act as an agent of nitrogen fixation using symbiosis with soil bacteria such as rhizobia, enabling them to efficiently convert atmospheric nitrogen into an available form in the soil to be used by the host and other crops^5^. Consequently, legumes are well suited for integration into low-input cropping systems, reducing the emissions of greenhouse gases^2^. Furthermore, legumes are a valuable diversification crop in modern agricultural practices^2,6^. They participate in crop protection by disrupting the pests’ and diseases’ life cycles^7^. As a result, they play a crucial role in promoting the development of sustainable farming systems for healthy agriculture^2,3^.

Species of this diverse family face significant economic losses due to pests, particularly those that feed on seeds in storage areas or fields^8^. In response to such threats, legumes have developed various defense mechanisms to protect their seeds against insect attacks. These defense mechanisms include structural barriers, secondary metabolites, and anti-nutritional compounds^9^. Of these compounds, specific albumin proteins have exhibited potent toxicity against storage pests^10,11^. This discovery has paved the way for developing greener alternative methods to safeguard stored grains.

In legumes, specifically germinating seeds, Albumin 1b (A1b) proteins are synthesized and serve as the primary source of sulfur amino acids^12^. Among these proteins is Pea albumin 1b (PA1b). PA1b is a cysteine-rich peptide of 37 amino acids belonging to the cysteine-knot/Albumin I family^13^ characterized by a knottin domain. The structure reveals an inhibitor cystine knot-like (ICK) fold comprising 3 beta strands. This knot is obtained when one disulfide bridge crosses the macrocycle formed by 2 other disulfides and the interconnecting backbone. The knottin structure is found in some plant protease inhibitors, cyclotides, toxins from spiders, insects, scorpions, and some antimicrobial peptides^13^.

PA1b is a hormone-like peptide found in the seeds of the pea plant and involved in the signal transduction system to regulate plant growth and differentiation^14^, and have been shown to have insecticidal properties^10,15^. In fact, PA1b has shown toxic activity in weevils (*Sitophilus* sp.)^10^ as well as in other insect species^11,16^. The toxic activity of this peptide is through ingestion^15^. PA1b binds and inhibits the activity of V-ATPase, a proton pump necessary for the absorption of nutrients^17^.

PA1b is derived from the expression of the *PA1* gene present in peas (*Pisum sativum*)^18^. The *PA1* gene compromises two exons separated by a single intron. The gene encodes for a signaling peptide, the A1b peptide product and its propeptide, and the A1a peptide product and its propeptide. Upon gene transcription, mRNA is produced and subsequently translated into a PA1 preproprotein which houses the signal peptide. Through post-translational modifications, two mature peptides are formed: the N-terminal fragment known as “PA1b” (3.8k kDa; 37 amino acids) and the C-terminal fragment referred to as “PA1a” (6kD; 53 amino acids)^18^. The synthesis of these 2 peptides is present in the different PA1 isoforms coexisting in peas and in various species of the Fabaceae. In this context, the diversity of albumin 1b peptides has been investigated using various molecular, biochemical, and bioinformatics tools.^18-20^.

The conserved structure and bioactivity of PA1b within the Fabaceae family led to the discovery of 6 isoforms of PA1b in pea (*Pisum sativum*) underlying the multigenic character of this plant entomotoxin^19^. Moreover, screening of 88 legume species across the three main legume subfamilies resulted in the identification of 19 novel *A1* genes from different tribes within the Faboideae subfamily but none from Caesalpiniodeae or Mimosoideae^21^. Although PA1b was not biochemically detected in barrel medics’ seeds (*Medicago* species), the genome analysis of the model legume *Medicago truncatula* (Trifoleae) revealed the presence of 53 homologous genes. Additionally, 21 copies were found in *Phaseolus vulgaris* (Phaseoleae), 7 copies in *Cajanus cajan* (Phaseoleae), and 3 copies in *Glycine max* (Phaseoleae)^22^. A PA1b sequence was also characterized in the seeds of *Lens culinaris* (Fabaeae)^23^ and in the roots of *Astragalus membranaceous* (Galegeae)^24^. So far, A1b homologs have been mainly uncovered from different species of the Faboideae subfamily of the legume family. Remarkably, copies of PA1b-like were found in species of the parasitic plant of the genus *Phelipanche* and *Orobanche* (family Orobanchaceae), which probably had the opportunity to have albumin 1 sequence through horizontal gene transfer due to their parasitic mode of life.

The biological activity of various legume seed flour^19^, seed extract, and seed peptide (PA1b-like) have been evaluated on both PA1b-susceptible and resistant strains of weevils^19,21^ and varied from none to highly acute depending on the concentration of PA1b found in the flour, the presence of other defense compounds in the seed, the type of solvent used for seed extraction, and the type of PA1b isoform isolated from the seed^19,21^. Additionally, chemically synthesized homologs of PA1b derived from *Medicago truncatula*, have been studied to assess their affinity for the binding site as well as their lethal concentration on cultured *Spodoptera frugiperda* Sf9 cells^22^. The results demonstrated that variants exhibiting a toxic effect showed different binding affinity and toxicity according to proper folding of the peptide^22^ and the presence of conserved key residues specific for the recognition and the interaction of PA1b via the membrane protein-based receptor sensitive to PA1b found in various insect species and responsible for its activity^25^. This lead to the discovery of a highly toxic peptide named AG41, exhibiting ten times greater toxicity compared to PA1b and the study of its structure and biological activity^22,26^.

This study aims to investigate the diversity of Albumin 1b (A1b) homologs in legumes native to the Middle East (ME) region, focusing on the Faboideae subfamily within the Fabaceae family. Specifically, degenerate primers were designed for the detection and genomic characterization of A1b homologs. The selected seeds from the ME region underwent sequence structure analysis of A1b variants, and phylogenetic relationships were explored among identified A1b sequences and publicly available A1b homologs in databases.

The study examines patterns of conservation and diversification patterns among A1b homologs of the selected species, consistent with the divergence of lineages within this plant group. In addition, the research assesses the toxicity of seed flour obtained from selected legume species in the ME region. In particular, variability in toxicity will be observed both between different species and within the same species from different geographical locations. Overall, the aim is to improve our understanding of A1b homologs in Middle Eastern legumes, their structural diversity, and potential implications for pest resistance in agricultural practice.

## Materials and methods

### Plant Material

Legume seeds from different Middle Eastern origins were obtained from the genetic resources collection of the International Center for Agricultural Research in the Dry Areas (ICARDA). Mainly, seeds belonging to the Faboideae (=Papilionoideae) subfamily of the Fabaceae (legumes) family were selected for this study. The collected seeds included species from various tribes, such as *Vicia, Lens*, and *Pisum* from the Fabeae tribe, *Medicago*, and *Astragalus* from the Galegeae tribe. **Table 1** shows all the tested species and their corresponding IG numbers assigned by ICARDA.

**Table 1.**
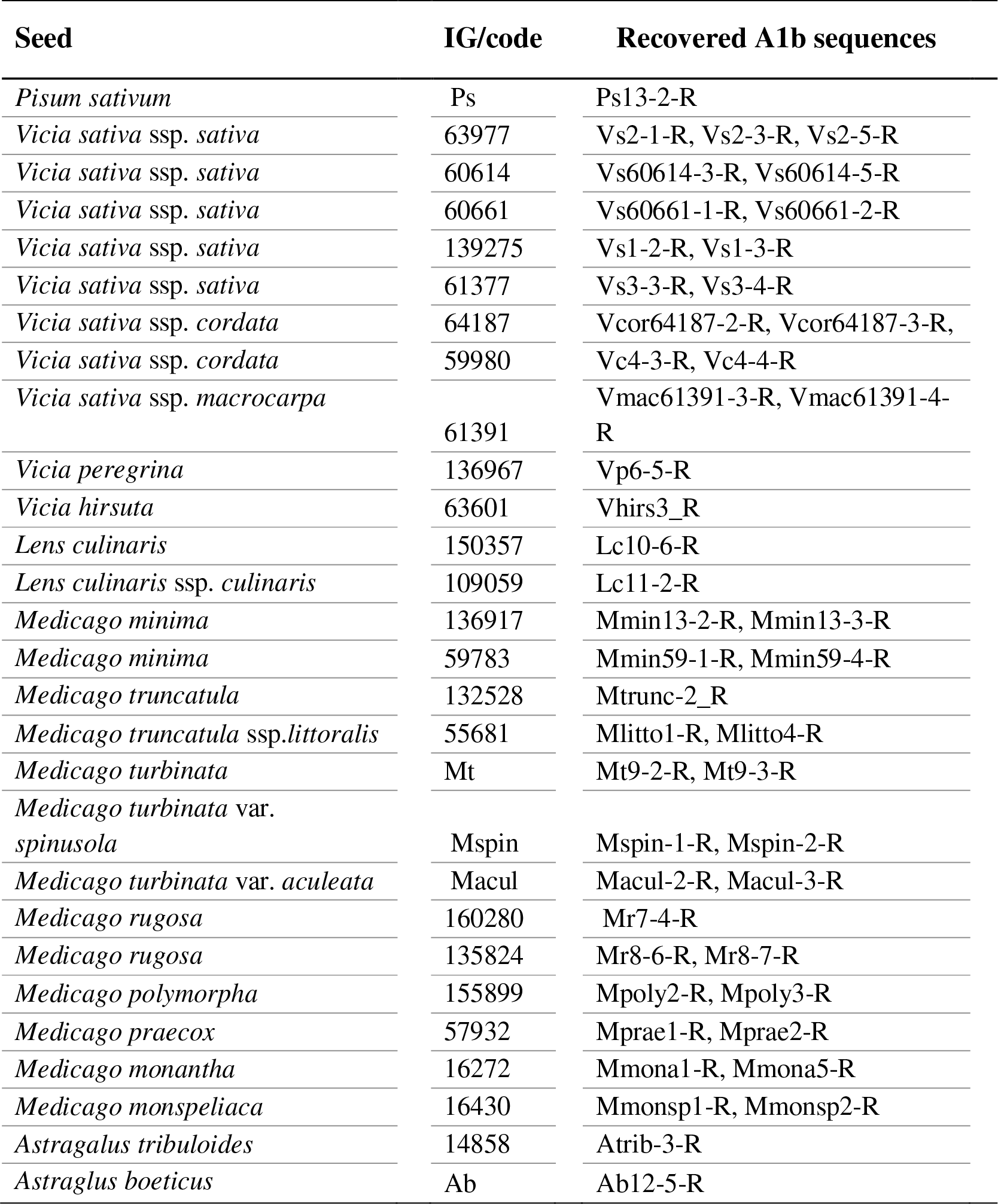
A1b sequences detected in each species from different origins through genomic characterization. The code given for each species is used for further sequence analysis.

### In vivo A1b Activity

#### Weevils

In this study, two strains of rice weevils (*Sitophilus oryzae*, Coleoptera, Curculionidae) were used. The susceptible strain, referred to as “WAA42,” was reared on wheat seeds (*Triticum aestivum*). On the other hand, the PA1b resistant strain, referred to as “ISO3R2”, housing the recessive pea-resistance allele, was reared on split peas (*Pisum sativum*). Weevils were maintained under controlled conditions of 27.5 ± 0.5 ° C and 70 ± 0.5 % relative humidity in a ventilated incubator (Memmert) kept in the dark.

#### Toxicity bioassays

Toxicity bioassays were conducted on both strains of weevils. As described by Delobel *et al*.,^10^ each bioassay involved placing 30 adult weevils, aged two weeks, on wheat-based flour dumplings containing the desired seed flour. Whole flours were tested at a concentration of 20% (w/w in wheat), compromising 50mg of seed flour, 200mg of wheat flour, and 160μl of ultrapure water. Wheat flour-based bait was used as a negative control, and Pea flour-based bait was used as a positive control. The prepared dumplings were left to dry overnight and then placed in cylindrical grilled plastic boxes measuring 2 cm in diameter and 4cm in height. The insects were incubated with the food source, and mortality was monitored for up to 14 days. The data obtained were analyzed by survival fitting analysis using the free software Simfit (http://www.simfit.man.ac.uk), where LT50s (lethal time 50%) were calculated.

### A1b homology

#### DNA Extraction

Seeds were ground to a fine powder for DNA extraction. A few seeds from each species were placed in an Eppendorf tube and mechanically crushed using a tissue lyser (Qiagen, Hilden, Germany) using two stainless steel beads (2.4 mm). The dried powder of each plant species was stored overnight at -80°C prior to DNA extraction. DNA from 40 mg of fine powder of each *Vicia* species, *Lens*, and *Pisum sativum* was extracted using DNeasy Plant Mini kit (Qiagen, Hilden, Germany) following the manufacturer’s instructions. However, modifications were made to extract DNA from 10 mg of each of the *Medicago*, and *Astragalus* species. Modifications were implemented after several trials to improve DNA quality and maximize yield. In the lysis step, a micro pestle was used to ease the lysis process and avoid any visible tissue clumps. In addition, the incubation of plant material with the lysis buffer was increased to 20 min. In the elution step, the DNA was not passed through the column as per the standard protocol. Instead, precipitation was performed by centrifuging the DNA for 15 minutes at maximum speed. The DNA was washed with 70% ethanol and centrifuged for 5 minutes at 8000 rpm. After air drying for 15 minutes, the DNA was resuspended in 100 μl TE buffer supplied by the kit. The concentration of DNA was measured using a NanoDrop spectrophotometer (ThermoFisher Scientific, Waltham, MA, USA), and the quality of DNA was assessed on 1% agarose gel electrophoresis in 1x TAE (Tris-Acetate-EDTA) buffer (Thermofisher Scientific).

#### Genomic amplification with degenerate primers

PCR was performed in a total of 50 μl reaction mixture containing 200 ng of genomic DNA and the Taq’Ozyme polymerase (1000 units, Ozyme) according to the manufacturer’s specifications with the following degenerate primers: Fcod forward: 5’-GYWMDGGKGYTTGTTCDCCDTTTG-3’ and R_2_ex_2_ reverse: 5’-AAACACCAKCCRTRKTCRAT-3’ (final concentration of 1 μ M each). These primers were designed based on aligned *PA1* like sequences from *Pisum sativum, Phaseolus vulgaris, Glycine max*, and *Medicago truncatula* found in the NCBI database to amplify a fragment of *PA1* gene from 400 to 600 bp according to the length of the gene coding for the signal peptide, PA1b and a part of PA1a sequence. Amplifications for PCR were carried out on a T100 thermal cycler (Biorad, USA) with the following cyclic parameters: initial denaturation at 94 °C for 1 min, 35 reaction cycles of 94 °C for 1 min (denaturation), 52 °C for 45 s (annealing) and 72 °C for 45 s (extension), and at 72 °C for 5 min for the final extension. Aliquots of PCR products were loaded with 1x Orange G loading dye (Thermo Fisher Scientific) and assessed by 1x TAE (Tris-Acetate-EDTA) agarose gel electrophoresis on a 1% (w/v) TAE prestained with 1 μl/10 ml GelRed (Biotium, Hayward, CA) at 100 V for 30 min. The ready-to-load DNA ladder mix (Euromedex) was used with a DNA marker size range between 80 bp and 1000 bp. Gels were viewed on an ultraviolet transilluminator and photographed with the Molecular imager system Gel Doc 1000 (Bio-Rad Laboratories, USA) using ImageLab software (BioRad, California).

#### Cloning

PCR amplified fragments were purified using a GenElute PCR Clean-Up kit (SIGMA). Purified PCR products were ligated to TOPO-TA vector pCR2.1 TOPO (Invitrogen, Carlsbad, CA, USA, catalog number 450641) and then transformed into chemically competent *Escherichia. coli* Top10 cells (Invitrogen) according to the supplier’s recommendations. Ten independent positive colonies were cultured in 2 ml Luria broth LB medium for each sample with 50 μg/mL of ampicillin. Tubes were placed on a Thermoshaker overnight at 37 °C. Following the protocol provided, five bacterial pellets containing plasmid DNA for each sample were then isolated using the Nucleopsin Plasmid-Plasmid DNA purification kit (Macherey Nagel). The other five bacterial pellets were stored at - 20°C. Isolated plasmids were then analyzed by restriction analysis (EcoRI restriction enzyme, Roche), and interesting bands were selected for sequencing.

### DNA Sequencing and Sequence Analysis

Plasmis DNAs were sequenced from both ends using Sanger sequencing carried out by Biofidal (Villeurbanne, France). The resulting sequences were cleaned to obtain the amplified fragment and then translated into the amino acid sequence Subsequently, the genomic sequence was compared with the reference sequence of the *PA1* gene from *Pisum sativum* (Genbank accession number: M13709). This comparison allowed for the description of the topology of the gene, including the identification of exon 1 coding for the signal peptide, intron, exon 2 coding for another part of the signal peptide, and the mature peptides PA1b and PA1a. The amino acids generated by the intron sequence were removed to refine the selected translated sequence. This was achieved by comparing the translated sequence with the protein reference sequence from *Pisum sativum* (AAA33638.1). The final defined pre-pro-proteins of all A1b-like sequences were further analyzed for the presence of potential signal peptide cleavage sites using the SignalP5.0 program (https://services.healthtech.dtu.dk/services/SignalP-5.0/) and all protein features were finally described. Therefore, our study used partial A1 like sequences from *Pisum, Vicia, Medicago*, and *Astragalus* (**Table1**), previously uncovered partial A1 like sequences using homology-based genome amplification from *Lablab purpureus* (LABPU), *Astragalus zachlensis* (ASTZA), *Astragalus angulosus* (ASTAN), *Astragalus ehdenensis* (ASTEH), *Astragalus trichopterus* (ASTTR), and *Medicago rotata* (MEDRO) found in the Middle East region (Karaki, 2013), complete A1 sequences from *Medicago truncatula* (Medtr8g022420: accession number XP_003627385.1, Medtr8g022400: XP_039685559.1, Medtr8g022430: XP_003627386.2, Medtr3g436100 (EG41): XP_013459315.1, TA24778_3880 (AG41): XP_013459315.1, Medtr6g017170 (DS37): XP_013451278.1, Medtr6g017150 (AS37): XP_013451275.1) from a previous study (Karaki *et al*., 2016) and different A1 sequences within legume species whose full genome sequencing and assembly has been completed and publicly available: *Pisum sativum* (PeaAlbumin1E: P62930.1, PeaAlbumin1D: P62929.1, PeaAlbumin1A: P62926.1, PeaAlbumin1F: P62931.1, PeaAlbumin1B: P62927.1, PeaAlbumin1C: P62928.1); *Cajanus Cajan* (C.cajan_46537: XP_020208801.1, C.cajan_40957: XP_020204276.1, C.cajan_39748: KYP36216.1, C.cajan_44236: XP_020208148.1); *Glycine max* (Glyma13g26330, Glyma13g26340); and *Phaseolus vulgaris* (Phvul011G205200: XP_007133739.1, Phvul011G204600: XP_007133733.1, Phvul011G205600: XP_007133743.1, Phvul011G205300: XP_007133740.1, Phvul011G203800: XP_007133725.1, Phvul011G203700: XP_007133724.1, Phvul011G204700: XP_007133734.1, Phvul011G205000: XP_007133737.1, Phvul011G203900: XP_007133726.1, Phvul011G205100: XP_007133738.1, Phvul011G204100: XP_007133728.1, Phvul011G204200: XP_007133729.1).

### Multiple alignment and Phylogenetic analysis

PA1b homologs from legume species gathered from this study were aligned using CLUSTAL OMEGA. The alignment was carefully inspected and manually adjusted using SeaView v5.0.5^27^. The phylogenetic tree of the protein sequences was inferred using the PhyML algorithm implemented in Seaview v5.05 program. The Le and Gascuel^28^ model was selected for the maximum likelihood inference using the “optimization” option of seaview to choose the invariable sites numbers and the variation rate across the different sites. The Best tree from the Nearest Neighbor Interchange (NNI) and Subtree Pruning and Regrafting (SPR) methods was chosen. To assess the robustness of the tree, a parametric bootstrap procedure was conducted with 500 replicates of the original datase^27^.

*Styphnolobium japonicum*, whose genome has been completely sequenced (Genbank accession number: GCA_023212625.1), is the ancestral species of our collection if we refer to the phylogeny of the Faboideae^29^. As the 8 sequences homologous to PA1b that we have detected in this work in the genome of *S. japonicum* systematically grouped together in all the phylogenetic trees that we have produced, we choose to use these sequences to root our phylogenetic tree.

## Results and Discussion

### Homology-based amplification of A1 sequences

Amplification of partial *A1* gene homologs was performed in 15 Faboideae species from different Middle Eastern origins using degenerate primers as described in the Materials and Methods section. **Supplementary file 1** displays the results of genomic DNA amplification from all tested seeds. As expected, bands of the appropriate size were obtained from selected species. *Pisum, Vicia*, and *Lens* species revealed interesting A1 bands ranging from 400 to 500 base pairs (bp) among the tested species. *Medicago* and *Astragalus* species showed bands between 400 and 600 bp.

In our study, a total of 81 new sequences from 15 different legume species from *Pisum, Vicia, Lens, Medicago*, and *Astragalus* of the Faboideae subfamily were obtained through cloning and sequencing. Following translation of each sequence in the six open reading frames and subsequent analysis of sequence topology, 61 sequences displayed the characteristic topology of the Albumin 1 family. Among these 61 sequences belonging to the A1 family, 13 sequences were found to be identical within the same species and origin. These sequences were determined to be duplicated and were subsequently excluded from further analyses. As a result, a total of 48 unique sequences from each species of different origin remained for subsequent analyses and multiple sequence alignment. **Table 1** provides an overview of the different amplified sequences corresponding to A1b variants detected in each species of different origins. The table highlights the presence of multiple variants of A1b within the same species in different legume species.

### Structure of A1b homologs

All A1b homologs were predicted to contain a signal sequence, an A1b sequence coding for Albumin 1 subunit b followed by the first propeptide, a partial A1a sequence coding for Albumin 1 subunit a with its propeptide (**Fig. 1**).

**Figure 1.**
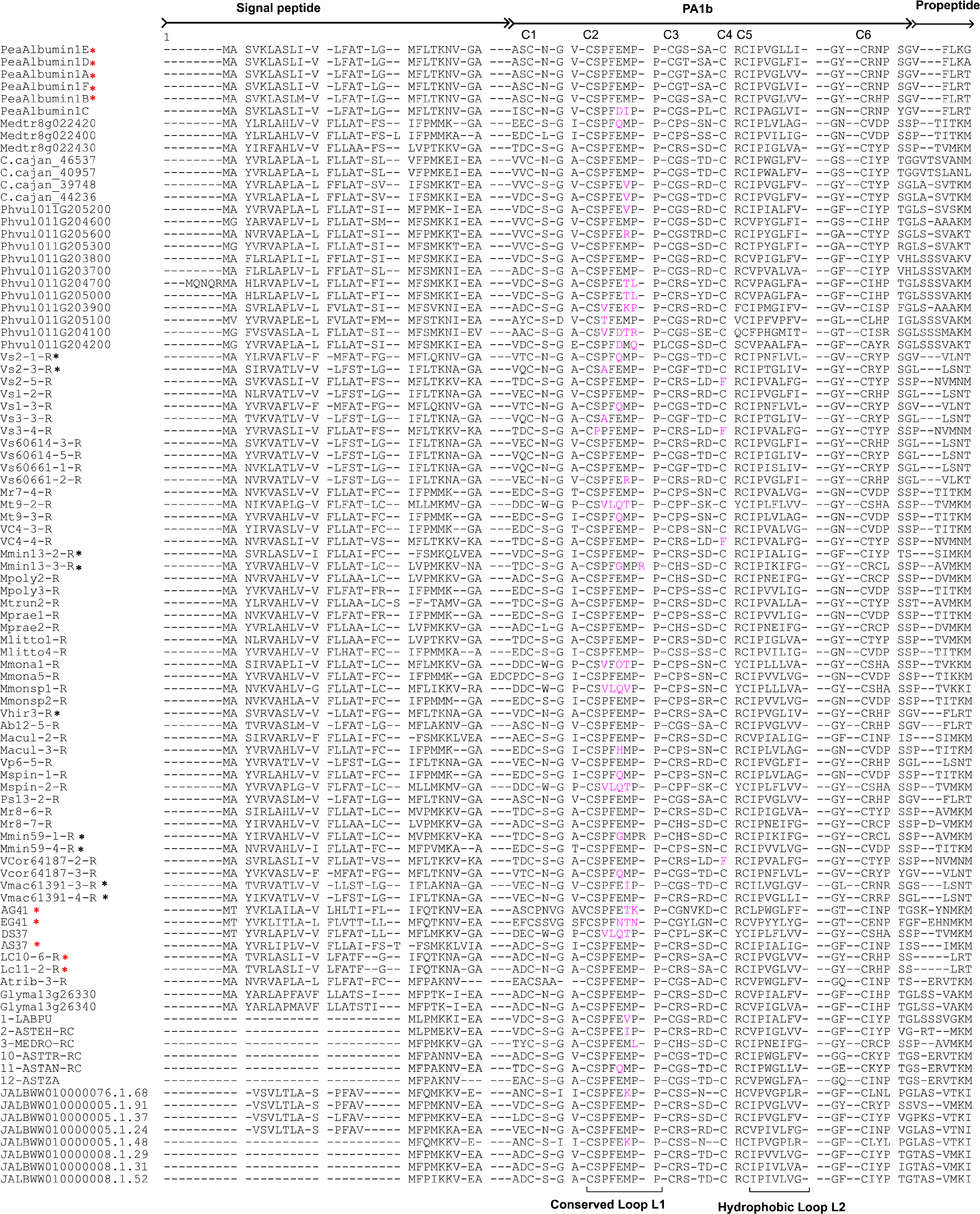
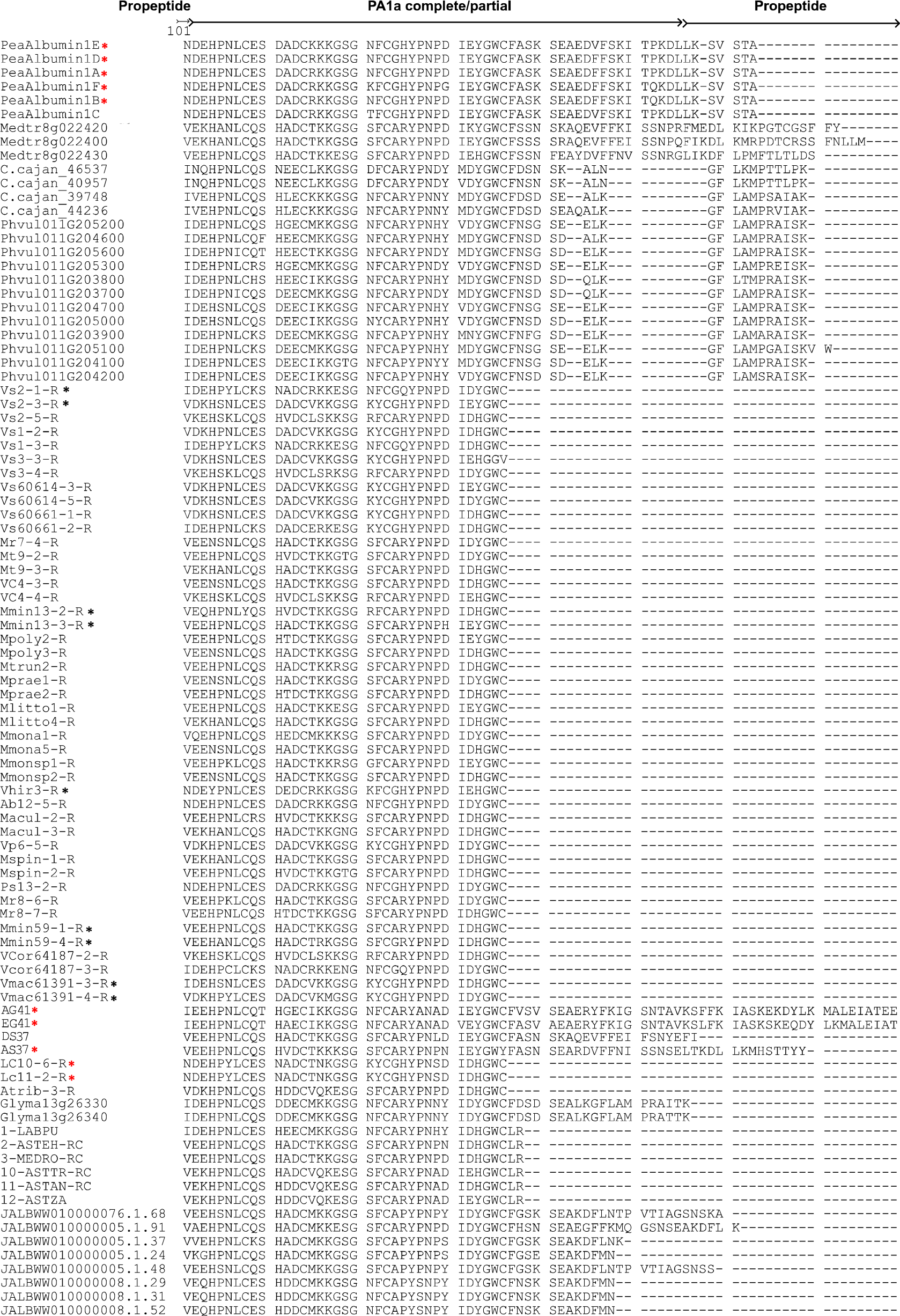
Protein alignment of partial cloned A1 sequences from selected legume species native to the Middle East region and characterized by our molecular approach (refer to Table 1).Additional sequences are used which are previously characterized partial A1 sequences (MEDRO: *Medicago rotata*, LABPU: *Lablab purpureus*, ASTZA: *Astragalus zachlensis*, ASTAN: *Astragalus angulosus*, ASTEH: *Astragalus ehdenensis*, ASTTR: *Astragalus trichopterus)*, reference PA1 sequences from *Pisum sativum*, and reference sequences from genomes of *Phaseolus vulgaris, Cajanus Cajan, Glycine max*, and *Medicago truncatula* publicly available. Variations in the key amino acids of the conserved loop L1 are highlighted in pink. Phenylalanine instead of cysteine 5 in some sequences is also highlighted in pink. A black asterisk corresponds to the putative A1b sequences that displayed toxic in vivo activity similar to or higher than PA1b isoforms found in peas. A red asterisk indicates the A1b sequences that are known to exhibit insecticidal activity.

We followed the annotation of *Pisum sativum* A1 preproprotein including PA1b sequence (37 amino acids) and PA1a sequence (53 amino acids). Three disulfide bridges engage 3 cystine binding C1-C4, C2-C5, and C3-C6 at positions (3-20), (7-22) and (15-32) respectively. Moreover, a triple-stranded antiparallel beta-sheet (residues 4 to 7, 21 to 23 and 30 to 33) along with the disulfide bonds constitute a signature of the ICK family. Furthermore, the L1 loop, composed of 8 residues (CSPFEMPP), through which the third disulfide bridge threads, is highly conserved in the PA1b structure. Finally, the hydrophobic loop L2 exposes hydrophobic residues at the molecular surface that are well conserved among PA1b homologs.

In our study, known and predicted A1b-like peptides length varied between 37 and 41 amino acids. However, PA1b characteristic sequence features are conserved in all PA1b homologous sequences as shown in the alignment where most residues are well conserved (**Fig. 1**).

In this context, L1 loop residues are conserved in the recovered sequences. However, different patterns were observed. *Pisum sativum* homologs show two residue variations Pea Albumin 1C (CSPFDIPP). In *Medicago* sp. and *Vicia* sp., variability is detected in five residues showing the following topologies: CS (P/V) (F/L) (E/Q/G/N) (M/V/T) (P/L/K) P and C (S/P) (P/A) F (E/Q) (M/I) PP) respectively. As for known sequences from databases, the following variations are detected in *Phaseolus vulgaris* and *Cajanus cajan* sequences respectively: CS (P/V/T) F (E/D) (M/T/R/V) (P/R/L/Q) P and CSPFE (M/V) PP (**Fig. 1**).

All sequences contained hydrophobic residues in the L2 loop and a high sequence similarity in the region of the beta sheet. More precisely, all sequences characterized by our molecular biology approach conserved six cysteine residues implicated in the disulfide pairing. However, C4 was missing in the following sequences Vcor64187-2-R, Vc4-4-R, Vs3-4-R, and Vs2-5-R where a Phenylalanine (F) was present instead of C4 resulting in 5 cysteine residues, and thus a loss of the cystine knot (**Fig. 1**).

The signal sequence varied in length from 25 to 28 residues in aligned sequences, potentially leading the mature protein through the secretory/protein body pathway. A higher conservation of residues was observed in the A1a sequences compared to A1b (**Fig. 1**).

### Phylogenetic analysis of A1b homologs

This study analyzed sequences of A1b homologs from legumes within the Faboideae subfamily. Overall, 48 A1b protein predicted sequences obtained after genomic amplification from seeds of different species from ME origin (**Table 1**), 6 sequences recovered by homology-based PCR from a previous study^22^, 7 sequences from *Medicago truncatula* previously characterized, 8 sequences recovered from the genome of *Styphnolobium japonicum* using BLAST tool, and 24 sequences from A1b homologs representatives of the Fabaceae family available publicly (**Fig. 2**) were aligned and used to draw the Maximum Likelihood phylogenetic tree.

**Figure 2.**
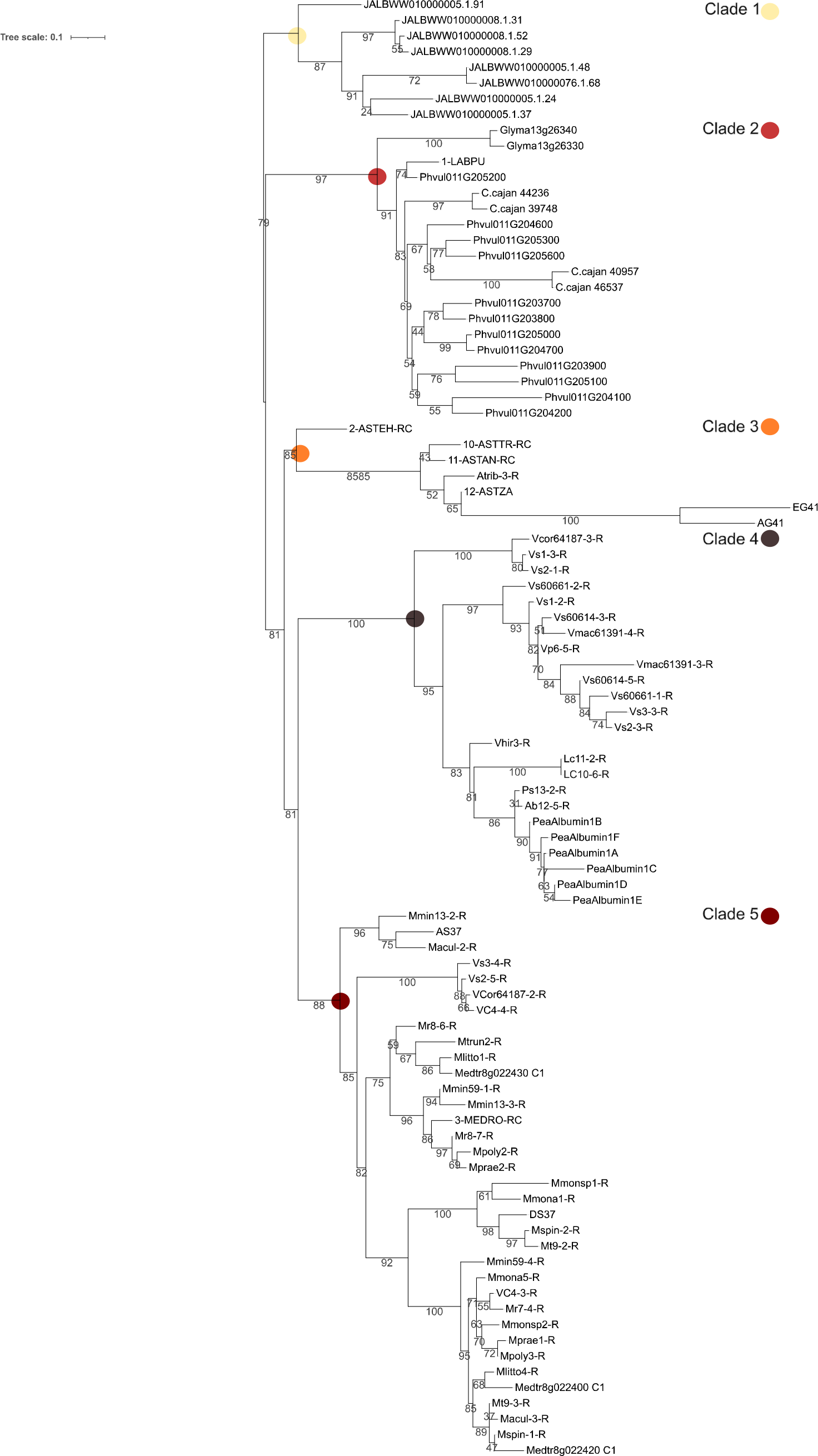
Maximum likelihood (LG model) phylogenetic tree of 93 A1b protein sequences from selected legume species native to the Middle East region. Bootstrap values expressed as percent (out of 500 replicates) are shown when >30%. Refer to Table 1 for the coding of the sequences realized by our PCR homology. Reference sequences characterized previously through a molecular approach are the following (MEDRO: *Medicago rotata*, LABPU: *Lablab purpureus*, ASTZA: *Astragalus zachlensis*, ASTAN: *Astragalus angulosus*, ASTEH: *Astragalus ehdenensis*, ASTTR: *Astragalus trichopterus*) Reference PA1b peptides from *Pisum sativum* are referred to as Pea Albumin (letter). Reference sequences from genomes of *Phaseolus vulgaris* (Phvul011), *Cajanus Cajan* (C. cajan), *Glycine max* (Glyma), and *Medicago truncatula* (Medtr) are used in the analysis. Reference sequences characterized from the genome of *Styphnolobium japonicum* are referred to as (JALBWW01000000.number).

The tree (**Fig. 2**) reveals 5 well-supported clades (Bootstrap values ≥85) containing A1b homologs of *Styphnolobium japonicum, Glycine max, Cajanus cajan, Phaseolusvulgaris, Lablab purpureus, Lens culinaris, Astragalus* sp., *Medicago* sp., *Vicia* sp., and *Pisum sativum*. The tree was rooted with *S. japonicum* sequences (**Fig. 2**). Phylogenetic analyses resolved the early diverging clades at the base of the Faboideae subfamily demonstrating that the Cladrastis clade containing *Styphnolobium* sp. among other clades are recognized as transitional between subfamilies Caesalpinioidae and Faboideae.

A1b sequences of *P. vulgaris, C. cajan, G. max* and *L. purpureus* are more closely related to each other than to any other A1b sequence from the study, they are grouped in clade 2. Furthermore, *P. vulgaris* and *C. cajan* sequences are more closely related to each other than to other sequences of clade2. This well-supported monophylectic group (Bootstrap 97) is confirmed in agreement with previous molecular phylogenetic studies^29-31^, it is called the Milletioid Clade and includes several legumes that are adapted to tropical climates, therefore, were named as warm season legumes, including *C. cajan, G. max*, and *P. vulgaris* as well as other legume species like *Vigna* sp. within the Faboideae subfamily.

A1b sequences from *Astragalus* sp. are clustered together with two sequences from *M. truncatula* (Clade 3) and are closely related to sequences from *Vicia* sp. (Clade 4) and *Medicago* sp. (Clade 5). The largest flowering plant *Astragalus* sp., as well as the most agriculturally important culinary pulses such as common beans, peas, lentils and soybeans are grouped in the Hologalegina Clade which is known to be a sister group to the Milletioid Clade and is divided into 2 subclades: Robinioids represented by *Lotus japonicus* and the Inverted Repeat-Lacking Clade (IRLC) which includes Cicer sp., *Lens culniaris, Medicago sativa, Vicia faba* and *Pisum sativum*. Finally, within the IRLC (Vicioid) subclade, phylogenetic studies supported a monophyletic group based on *Medicago* and *Trifolium* as sister to *Vicia*^29-31^. Sequences grouped in Clade 4 show sequences from *Vicia* sp. closely related to *Lens culinaris* and *Pisum sativum* sequences. However, few sequences from *Vicia* sp. are grouped together with sequences from *Medicago* sp. in Clade 5. These results indicate that 2 major duplication events occurred after the divergence of the Milletioid and the Hologalegina subclades leading to diversification of the A1b sequences in the evolution of different species from these 2 subclades, thus conserving major sequence structure motifs related to the PA1b function.

### Variability of seed flour toxicity

Seeds from species showing A1b gene/peptide structure were tested for biological activity on weevil strains. The toxicity of seed flour against both the PA1b-susceptible “WAA42” strain and the resistant “ISO3R2” strain of weevils varied significantly among legume species of different origins and even within the same species of different origins. Legume seeds are known to accumulate many compounds with insecticidal activity including PA1b molecules^10,32,33^. However, the variability in the toxicity of legume seeds is related to the biological activity of one or multiple compounds present in the flour assuming that A1b-like activities affect differentially the 2 weevil strains. **Table 2** provides the LT50% values (days), representing the survival time in days for 50% of the tested weevils’ populations for determination of the insecticidal activity of each seed flour tested on weevil strains. The results indicate that certain seed flours induced mortality in the susceptible strain of weevils but not in the resistant strain, suggesting a putative A1b activity. This activity was particularly prominent in species such as *Pisum sativum* (Ps) and *Vicia sativa* ssp. *sativa* (Vs63977), *Vicia sativa* ssp. *macrocarpa* (61391) [LT50 ≈ 8 days]; *Lens culinaris* (150357), *Lens culinaris* ssp. *culinaris* (109059), *Vicia hirsuta* (63601) [LT50 ≈ 6 days] as compared to the positive control, which is pea flour [LT50 ≈ 8 days] (*Pisum sativum* var. *Isard*) (**Table 2**). Only one species of *Medicago* out of 8 different species, namely *Medicago minima* (136917, 59783) [LT50 ≈ 5 days], displayed the highest toxicity on WAA42 strain but not on ISO3R2 strain, suggesting an A1b-like activity (**Table 2**). In other cases, toxicity was observed in both strains of weevils which suggests a possible synergistic effect between A1b-like peptides present in the seeds and other non-proteinaceous compounds found in legume seeds such as saponins^32^ (Da Silva *et al*., 2012). Alternatively, the observed toxicity could be attributed to the accumulation of certain compounds, such as polyphenols, tannins, and alkaloids^34^, found in legume seeds without A1b activity. Putative A1b activity in combination with other compounds was observed in species of *Medicago* and *Astragalus*, particularly in *Medicago turbinata, M. turbinata* var. *spinusola, M. turbinata* var. *aculeata, M. monspeliaca* (16430) [LT50 ≈ 6 days], and *Astragalus boeticus* [LT50 ≈ 5 days] (**Table 2**). This finding is interesting as other compounds present in the seed could be characterized and potentially used to control storage pests. On the other hand, some species of *Vicia* and *Medicago* showed minor or no toxicity toward both strains of weevils (**Table 2**) suggesting the absence of sufficient toxic compounds in the seeds including A1b-like molecules.

**Table 2.** Mean survival time (Lethal time 50%) of both susceptible and resistant strains of weevils (*S. oryzae*) on baits containing seed flour of the 28 selected seeds within the Faboideae subfamily native to the Middle East Region (15 different species). NE means No Effect. The IG number is assigned for the identification of legume species from the ME region. ^*^ Legume seeds showing an in vivo A1b-like toxic activity equivalent or higher than that of peas.

| Seed | IG/Code | Susceptible strain<br>"WAA42" LT50 +/-<br>SEM | Resistance strain<br>"ISO3R2" LT50<br>+/- SEM |
| --- | --- | --- | --- |
| T- Wheat flour |  | NE | NE |
| T+ Pois Isard |  | 7.6+/-0.4 | NE |
| *Pisum sativum* | Ps | 7.8 +/- 0.4 | NE |
| *Vicia sativa* ssp. *sativa* * | 63977 | 8.0+/-0.4 | NE |
| *Vicia sativa* ssp. *sativa* | 60614 | NE | NE |
| *Vicia sativa* ssp. *sativa* | 60661 | NE | NE |
| *Vicia sativa* ssp. *sativa* | 139275 | 14.4 +/- 0.5 | NE |
| *Vicia sativa* ssp. *sativa* | 61377 | 17.2+/- 2.7 | NE |
| *Vicia sativa* ssp. *cordata* | 64187 | NE | NE |
| *Vicia sativa* ssp. *cordata* | 59980 | 11.0+/-0.7 | NE |
| *Vicia sativa* ssp. *macrocarpa* * | 61391 | 8.06+/-0.612 | NE |
| *Vicia peregrina* | 136967 | 9.8+/-0.7 | NE |
| *Vicia hirsuta* * | 63601 | 5.79+/-0.28 | NE |
| *Lens culinaris* * | 150357 | 5.9 +/- 0.3 | NE |
| *Lens culinaris* ssp. *culinaris* * | 109059 | 6.3 +/- 0.4 | NE |
| *Medicago minima* * | 136917 | 5.58+/-0.26 | NE |
| *Medicago minima* * | 59783 | 4.33+/-0.2 | NE |
| *Medicago truncatula* | 132528 | NE | NE |
| *Medicago truncatula* ssp. *littoralis* | 55681 | 7.14 +/- 0.28 | 7.59 +/- 0.21 |
| *Medicago turbinata* | Mt | 4.8 +/- 0.1 | 6.4 +/- 0.5 |
| *Medicago turbinata* var. *spinosola* | Mspin | 5.85+/-0.25 | 6.6+/-0.28 |
| *Medicago turbinata* var. *aculeata* | Macul | 5.4 +/-0.26 | 4.6 +/-0.18 |
| *Medicago rugosa* | 160280 | NE | NE |
| *Medicago rugosa* | 135824 | NE | NE |
| *Medicago polymorpha* | 155899 | NE | NE |
| *Medicago praecox* | 57932 | NE | NE |
| *Medicago monantha* | 16272 | 9.35 +/-0.36 | 11.1 +/- 0.33 |
| *Medicago monspeliaca* | 16430 | 5.892+/-0.23 | 5.91 +/-0.31 |
| *Astragalus tribuloides* | 14858 | 12.2 +/-0.19 | 12.2 +/-0.19 |
| *Astragalus boeticus* | Ab | 4.7 +/- 0.4 | 5.0 +/- 0.6 |

In addition to PA1b isoforms known for their entomotoxic properties^35^, A1bs with known insecticidal activity from *Lens culinaris* (AHG94969.1)^23^ and *Medicago truncatula* including AG41 and EG41 as well as one identified A1b with a well-conserved gene structure and insecticidal activity, AS37, conserved CRC motif with its arginine residue, crucial for insecticidal activity^22^.

Partial A1 sequences from *Lens* species (Lc10-6-R and Lc11-2-R) native to the Middle East showed 91.3% identity with the partial insecticidal known sequence found in the database^23^ with 100% identity of the mature peptide A1b. Additionally, Vhir3-R from *Vicia hirsuta* 63601 showed 71% identity with the putative albumin 1 found publicly in *Vicia hirsuta* CAH05251.1^21^. The recovered sequence of our work Vhir3-R showed a unique segment (RSSA) after the 3^rd^ cysteine that is only found in the known insecticidal sequence from Lens (**Fig.1**).

The AG41 sequence displayed a very high toxicity, almost 10 times superior to that of the original PA1b^21^. The loop L2 contains the main hydrophobic residues that play a significant role in the binding and cellular toxicity of A1b peptides. The hydrophobic role in AG41 is formed by the following residues Leu27, Pro28, Trp29, Leu 31, Phe 32 and Phe 33 by facing residues Pro12 and Phe 13 of L1^26^, whereas PA1b displayed the following residues Val25, Leu27, Val28 and Ile29 with the facing residue Phe10 of L1^25^. Screening of hydrophobic residues in sequences from toxic seeds was done. In *V. hirsuta* (Vhir3-R), potential hydrophobic residues at the surface of the molecule show the following pattern (Val25, Leu27, Ile28 and Val29), whereas in *V. macrocarpa* (Vmac61391-3-R) and (Vmac61391-4-R) the following residues were displayed Val25, Leu27, and Val29 and Val25, Leu27, Leu28 and Ile29 respectively. *Lens* species (Lc10-6-R and Lc11-2-R) showed Val25, Leu27, Val28, and Val29. However, 2 out of 4 M. min (Mmin59-1-R, and Mmin13-3-R) displayed Phe28 in L2 which is also present in AG41 molecule. Moreover, all sequences from toxic seeds retained an arginine residue in the CXC motif (**Fig.1**).

Screening of other key residues required for binding and insecticidal activity of A1b-like molecules and present in every insecticidal PA1b-like peptide identified so far in legumes, consisted of 4 important residues (Phe10, Arg21, Ile23, and Leu27)^25^. All seeds from ME origin showing toxicity on susceptible weevils conserved these crucial residues.

## Conclusion

Our study recovered a total of 48 genomic A1 b-like sequences from *Pisum, Vicia, Medicago*, and *Astragalus* native to the Middle East region. The sequences showed variability of A1b within the same species and origin. However, they displayed the characteristic topology of the Albumin 1 family. These newly identified sequences were used with other A1b like sequences from Middle Eastern origin from a different study, as well as known A1b sequences from the database for phylogenetic analysis. Results show that genomic characterization and variability of recovered sequences follow species lineage positions. Moreover, the study of seed flour toxicity enabled us to identify novel candidates such as *Vicia sativa* and *Medicago minima* showing high A1b insecticidal activity against susceptible strains of weevils. Further investigation of the entomotoxic activity and molecular variability of A1b peptides from different legume species will provide important insights into the crucial A1b entomotoxic properties and will facilitate the development of enhanced A1 b-like insecticidal molecules for pest control.

## Supplementary file1

**Supplementary file 1:**
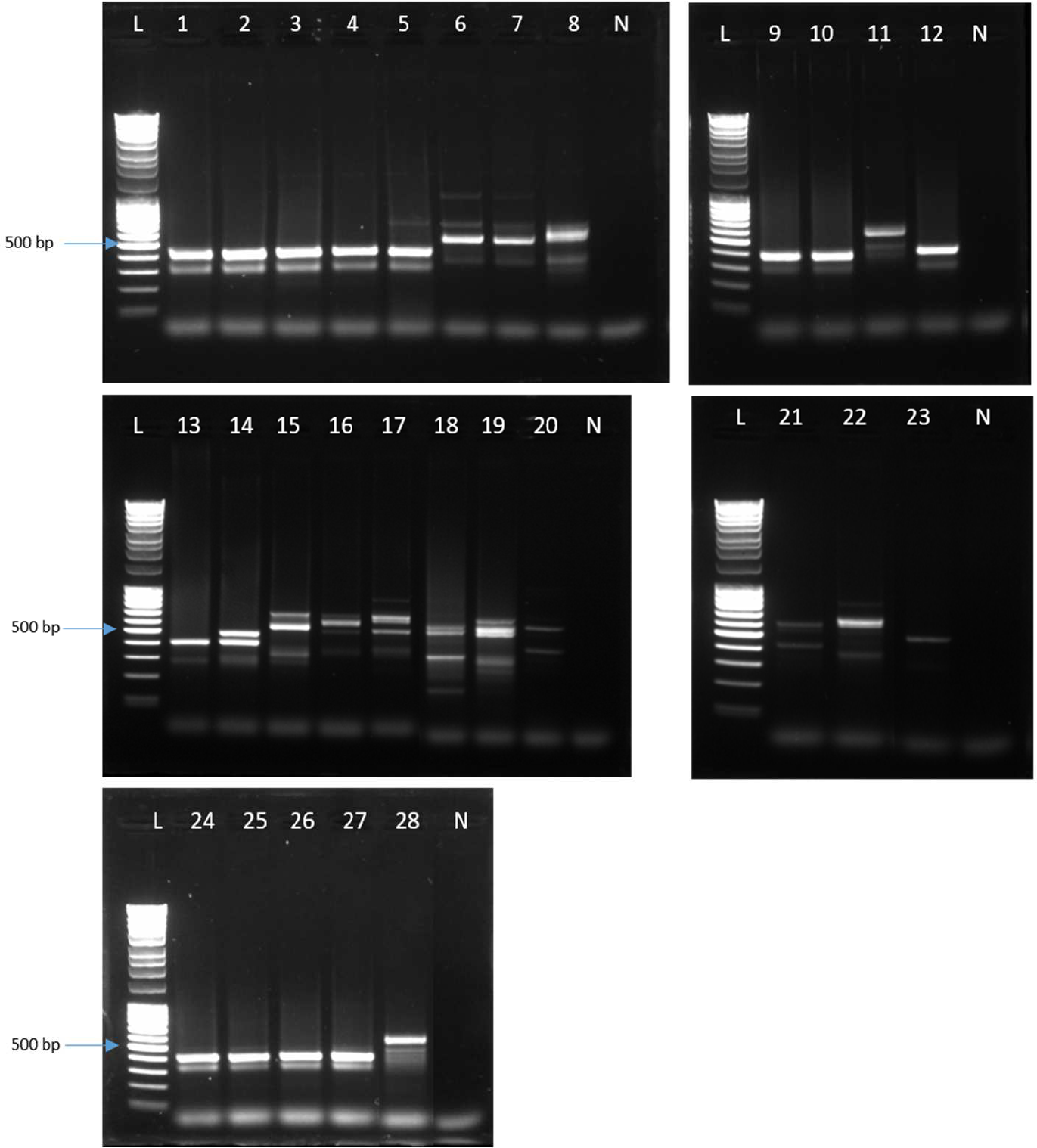
Agarose gel electrophoresis (1%) of DNA fragments of partial A1 gene amplified through PCR using degenerate pair of primer (Fcod, R_2_ex_2_) on genomic DNA of different legume species native to the Middle East. Lane L: Ladder euromedex 1kb; Lane N: Negative control; 1: *Vicia sativa* ssp. *sativa* (IG: 139275); 2: *V. sativa* ssp. *sativa* (63977); 3 *V. sativa* ssp. *sativa* (61377); 4: *V. sativa* ssp. *cordata* (59980); 5: *V. peregrina* (136968); 6: *Medicago rugosa* (160280); 7: *M. rugosa* (135824); 8: *M. turbinata*; 9: *Lens culinaris* (150357); 10: *L. culinaris* ssp. *culinaris* (109059); 11: *Astragalus boeitcus*; 12: *Pisum sativum*; 13: *V. hirsuta* (63601); 14: *M. truncatula* (132528); 15: *M. turbinata* var. *aculeata*; 16: *M. minima* (136917); 17: *M. praecox* (57932); 18: *M. truncatula* ssp. *littoralis* (55681); 19: *M. turbinata* var. *spinusola*; 20: *M. monatha* (16272); 21: *M. monspeliaca* (16430); 22: *M. polymorpha* (155899); 23: *Astragalus tribuloides* (14858); 24: *Vicia sativa* ssp. *sativa* (60614); 25: *Vicia sativa* ssp. *sativa* (60661); 26: *V. sativa* ssp. *cordata* (64187); 27: *V. sativa* ssp. *macrocarpa* (61391); 28: *M. minima* (59783)

